# PharmacoDB: an integrative database for mining *in vitro* drug screening studies

**DOI:** 10.1101/195149

**Authors:** Petr Smirnov, Victor Kofia, Alexander Maru, Mark Freeman, Chantal Ho, Nehme El-Hachem, George-Alexandru Adam, Wail Ba-alawi, Zhaleh Safikhani, Benjamin Haibe-Kains

## Abstract

Recent pharmacogenomic studies profiled large panels of cancer cell lines against hundreds of approved drugs and experimental chemical compounds. The overarching goal of these screens is to measure sensitivity of cell lines to chemical perturbation, correlate these measures to genomic features, and thereby develop novel predictors of drug response. However, leveraging this valuable data is challenging due to the lack of standards for annotating cell lines and chemical compounds, and quantifying drug response. Moreover, it has been recently shown that the complexity and complementarity of the experimental protocols used in the field result in high levels of technical and biological variation in the *in vitro* pharmacological profiles. There is therefore a need for new tools to facilitate rigorous comparison and integrative analysis of large-scale drug screening datasets. To address this issue, we have developed PharmacoDB (pharmacodb.pmgenomics.ca), a database integrating the largest pharmacogenomic studies published to date. Here, we describe how the curation of cell line and chemical compound identifiers maximizes the overlap between datasets and how users can leverage such data to compare and extract robust drug phenotypes. PharmacoDB provides a unique resource to mine a compendium of curated pharmacogenomic datasets that are otherwise disparate and difficult to integrate.

**Key points:**

- Curation of cell line and drug identifiers in the largest pharmacogenomic studies published to date
- Uniform processing of drug sensitivity data to reduce heterogeneity across studies
- Multiple drug response summary metrics enabling visual comparison and integrative analysis

## INTRODUCTION

Cancer has emerged as one of the principal causes of mortality in the 21st century (1). It is a collection of related diseases with widely different prognosis and response to therapy (2). This heterogeneity poses challenges for treatment, as patients with the same diagnosis often have different responses to treatment and may develop resistances at different rates (3). The genesis, progression, and response to pharmacotherapy of cancer is largely determined by the molecular state and features of the tumour cells (4). This observation spurred the development of the field of pharmacogenomics to study the relationships between genomic, transcriptomic and proteomic features of cancer cells and their response to treatment with small molecule compounds.

Immortalized cancer cell lines are the most widely-used models to study response of tumors to anticancer compounds (5). In addition to being comprehensively profiled at the molecular level, cancer cell lines can be cultured to conduct high-throughput drug screening studies, where large panels of compounds are screened for their efficacy of halting the growth or killing molecularly distinct cancer tumour models (6). Over the past decade, several large studies combining high-throughput *in vitro* drug screening with molecular profiling of cancer cell lines have been published (7–13). Recognizing that the molecular diversity of cancer cannot be faithfully represented by small panels of cell lines, these studies have assembled large panels of hundreds to over a thousand cell lines and profiled them at the molecular and pharmacological levels. These valuable data have been publicly released via well-established repositories, including NCBI Gene Expression Omnibus (14) and EMBL-EBI European Genome-phenome Archive (15), and institutional websites.

Recent computational approaches have leveraged these high-throughput pharmacogenomic data in a wide range of biomedical applications. Drug screening on hundreds of cell lines has enabled more comprehensive discovery of molecular features associated with drug response across cancer types and in specific tissues (16). Transcriptomic changes due to chemical perturbations in cell lines were intensively used to match drug to disease with the aim to define new indications for existing drugs, also referred to as drug repurposing (17, 18). Pharmacogenomic data have been used to develop new classification schemes for chemical compounds in order to determine or refine their mechanism of action (19, 20). *In vitro* drug screening data combined with single-cell RNA-sequencing profiles have opened new avenues of research for the rational discovery of synergistic drug combinations in renal cell carcinoma (21). Recent studies have investigated the relevance of *in vitro* and *in vivo* drug screening for precision medicine (22–24). These examples clearly demonstrate the value of the pharmacogenomic data for basic and translational cancer research.

The main limitation of the majority of published pharmacogenomic studies is that they are restricted to the analysis of single datasets. This is primarily due to inconsistent annotations of cell lines and compounds, which prevents direct comparison between datasets (25). Meta-analysis of pharmacogenomic data is further hindered by the lack of standards for statistical modeling of drug dose-response curves and subsequent summarization into drug sensitivity measures (25–28). However, joint analysis of independent datasets holds the potential to improve robustness of research outputs against variations in the complex experimental protocols used in high-throughput drug screening (29, 30). To address these issues we developed PharmacoDB, the first database integrating multiple high-throughput pharmacogenomic datasets that have been recently released(Table 1; Supplementary Figure 1A). PharmacoDB provides an intuitive interface to search and explore these datasets (Figure 1A) based on cell lines and their tissue source, compounds and their targets (Figure 1B), and experiments in which cell viability is measured for cell lines treated with chemical compounds (Figure 1C). Moreover, PharmacoDB provides access to molecular profiles of cell lines and computational analytical tools via linkage to *PharmacoGx* (Figure 1D; Supplementary Figure 1A), an R/Bioconductor package implementing a suite of statistical modeling functions to jointly analyze molecular features and drug dose-response curves (31). Here, we describe the content of our integrative pharmacogenomic database, the curation process, and its web-interface.

**Table 1.** Specifications of all the datasets included in PharmacoDB. Viability assay: Syto60: Proliferation; fluorescent DNA stain (Invitrogen); CellTiter Glo: Viability, membrane integrity, ATP (Promega); SRB: Sulforhodamine B colorimetric. Available molecular data: Mut: targeted mutation data; ExomeSeq: Whole exome-sequencing data; mRNA: gene expression data; methylation: methylation microarray data; CNV: copy number variation data; RPPA: protein expression data using reverse phase protein lysate microarray.

| Dataset | # Drugs | # Cells | # Experiments | Viability Assay | Available Molecular Data | Citations |
| --- | --- | --- | --- | --- | --- | --- |
| CCLE | 22 | 1,061 | 11,670 | CellTiter Glo | mRNA expression, CNV, Mutation | (9) |
| GDSC1000 | 250 | 1,109 | 225,480 | Syto60 | mRNA expression, CNV, Mutation, Methylation | (7, 8) |
| gCSI | 16 | 754 | 6,455 | CellTiter Glo | mRNA expression, CNV, Mutation | (11, 12) |
| GRAY | 90 | 84 | 9,413 | CellTiter Glo | mRNA expression, CNV, RPPA, Methylation, ExomeSeq | (10, 33) |
| FIMM | 52 | 50 | 2,561 | CellTiter Glo | None | (13) |
| CTRPv2 | 544 | 888 | 395,263 | CellTiter Glo | None | (19, 32) |
| UHNBreast | 4 | 84 | 52 | SRB | RNAseq, RPPA, Mut | (29) |
| **Total** | **759** | **1,691** | **650,894** |  |  |  |

**Figure 1.**
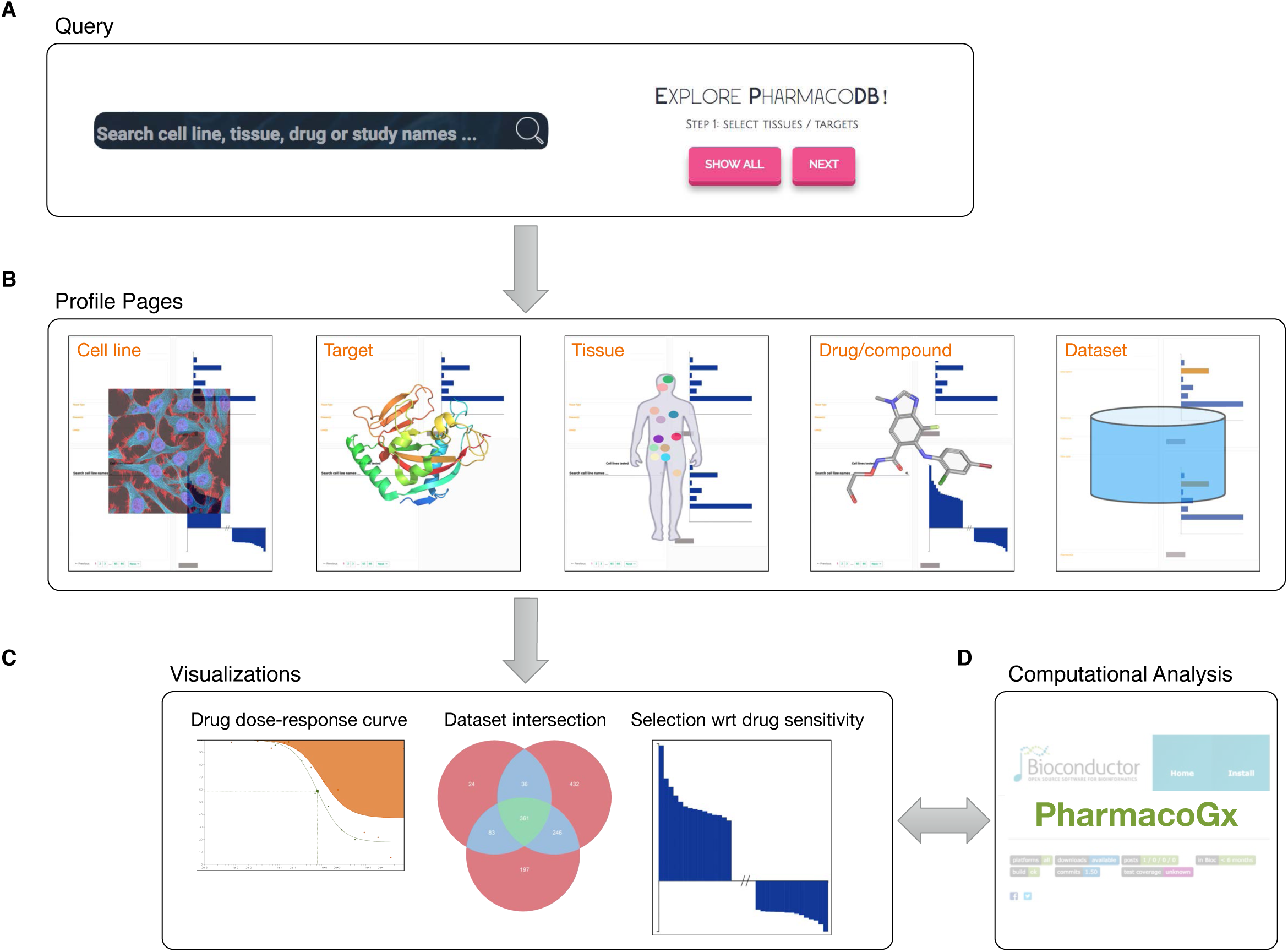
Main functionalities of PharmacoDB, displaying (**A)** the interfaces to query the database through searching or exploring available entities, (**B)** the five primary data types with respective profile pages, (**C**) the main visualizations of the aggregated data in PharmacoDB, and (**D**) the link to *PharmacoGx* for extensive computational analysis of pharmacogenomic data.

## DATA COLLECTION AND DATABASE CONTENT

### Pharmacogenomic studies

PharmacoDB seeks to include the largest published studies investigating the viability response of human cancer cell lines to chemical compound treatment. To date, we have curated 7 major studies: The Cancer Cell Line Encyclopedia (CCLE) (9), Genomics of Drug Sensitivity in Cancer (GDSC) (7, 8), Genentech Cell Screening Initiative (gCSI) (11, 12), the Cancer Therapeutic Response Portal (CTRP) (19, 32), the Oregon Health and Science University (OHSU) Breast Cancer Screen by Dr. Joe Gray’s lab (GRAY) (10, 33), the Institute for Molecular Medicine Finland cell viability screen (FIMM) (13), and the University Health Network (Toronto) breast cancer screen (UHNBreast) (29). For each study, we downloaded the cell line and compound annotations available with the original publications of the study, either through the journal website or dedicated portals for data sharing made available by the study authors (Table 1; Supplementary Figure 1A; Supplementary Methods).

### Annotation of cell lines and chemical compounds

We performed semi-automated curation of all the cell line and compound identifiers with the goal of discovering and maximizing the overlap between the datasets. First, we looked for exact case-insensitive matches of the identifiers used in the dataset undergoing curation to already curated unique identifiers, if applicable. Second, for all remaining compounds and cell lines, a partial matching algorithm was used to generate candidate unique identifier matches for each identifier used in the study. These candidate matches were manually reviewed to find the correct match for all compounds and cell lines which had a matching unique identifier. Third, for the subset of compounds for which there was no match using the compound names, we used any other provided compound annotations such as the SMILES, InchiKey or PubChem identifier to match them with identifiers available for previously curated compounds. Lastly, for compounds missing all of the SMILES, InchiKey and other chemical identifiers, the PubChem database was accessed through the *WebChem* R package (version 0.2) and queried by a compound’s name as described below. If it was possible to retrieve the identifiers of these compounds, then the third step was repeated to find possible matches with previously curated compounds. For cell lines which had no correct matches in the second step, Cellosaurus (34) was queried to generate candidate cell name synonyms, and matching was attempted with current unique identifiers. If at the end of these four steps there remained any cell line or compound names that were not matched, a new unique, human interpretable identifier was created based on the name from the dataset currently undergoing curation. This curation process maximized the overlap between datasets (Figure 2), and established links to external databases using Pubchem identifiers for compounds and Cellosaurus accession identifiers for cell lines. Overall, we identified 1,691 unique cell lines from 41 tissue sources (Figure 3A) and 759 unique compounds with 673 associated targets (Figure 3B).

**Figure 2.**
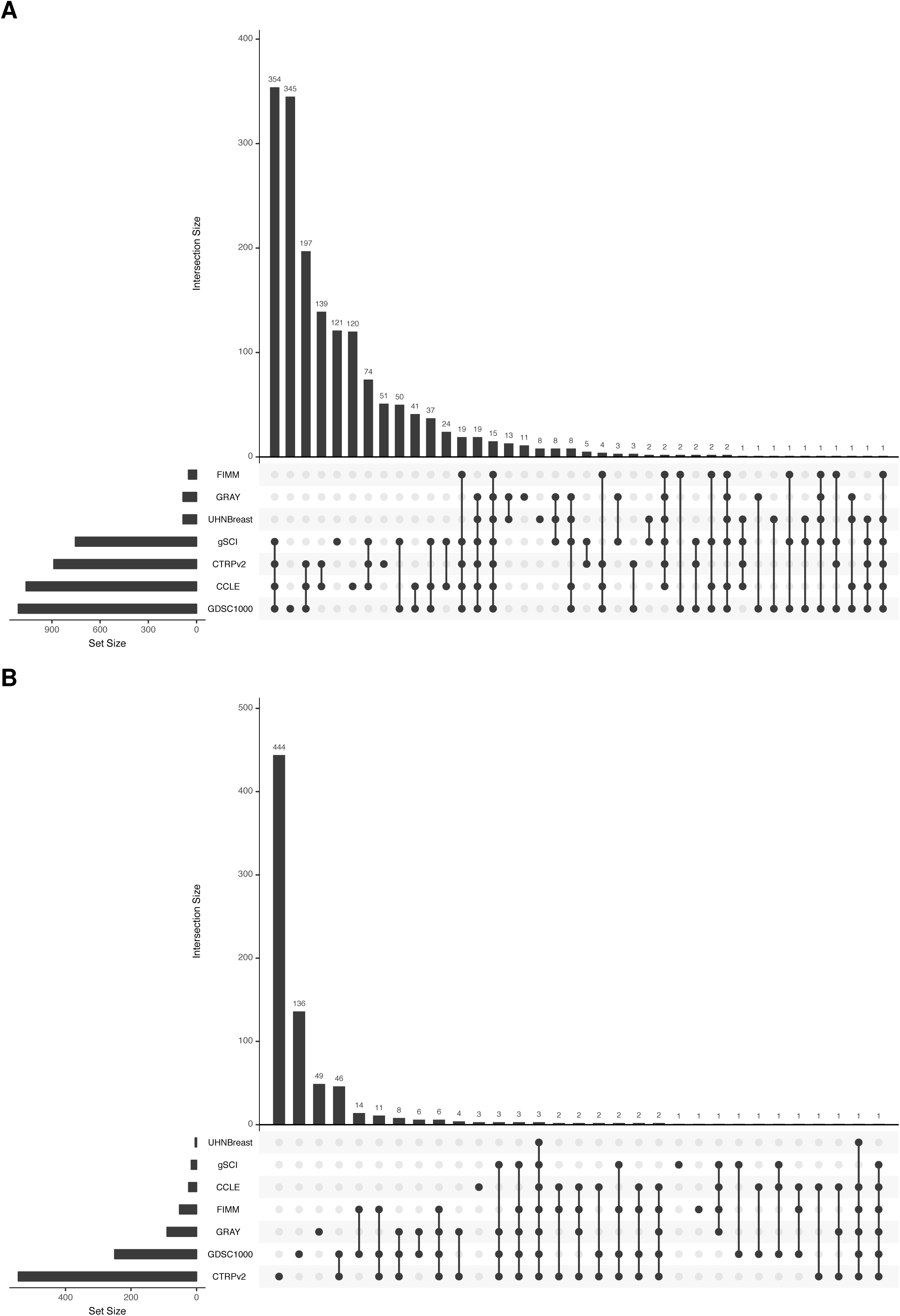
Intersection between datasets included in PharmacoDB for (**A**) cell lines and (**B**) compounds, after curation of cell line and compound identifiers, respectively.

**Figure 3.**
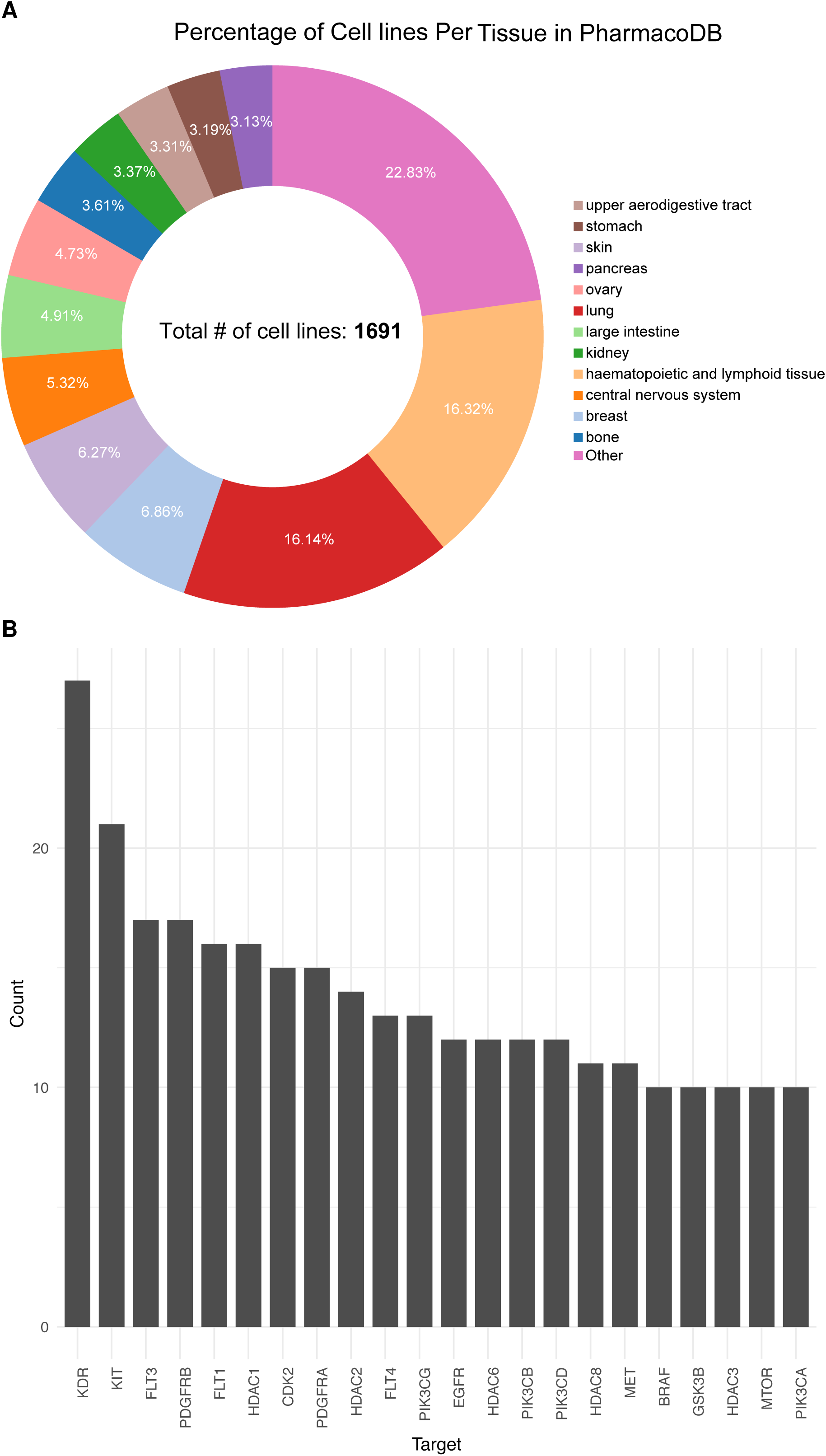
Cell lines and targets in PharmacoDB. (**A**) The proportion of cell lines per tissue type in PharmacoDB, after curation, and (**B**) drug targets in PharmacoDB annotated as targeted by 10 or more compounds.

Our curation process substantially increased the overlap across all the datasets compared to an exact matching procedure for both cell lines and compounds (Figure 4). While some intersections increased only modestly due to similar identifiers being used in the original studies, the benefit was substantial for others. For example, the intersection of compounds tested in GDSC1000 and CTRPv2 more than tripled, from 27 to 90 (Figure 4). While many of the newly matched identifiers differed only in capitalization or hyphenation, computational approaches to mapping identifiers which ignore these differences would be insufficient. These approaches would fail to match certain cases, such as the matching of compound names AZD6244 and Selumetinib, and would also cause mismatches, as for example for the distinct cell lines KMH-2 and KM-H2, which are respectively a Hodgkin’s lymphoma cell line and a thyroid gland carcinoma. Differences in naming conventions often create difficulties and confusion for researchers who wish to integrate data from across different studies, and the curation done in PharmacoDB aims to alleviate this barrier to leveraging these valuable pharmacogenomics studies.

**Figure 4.**
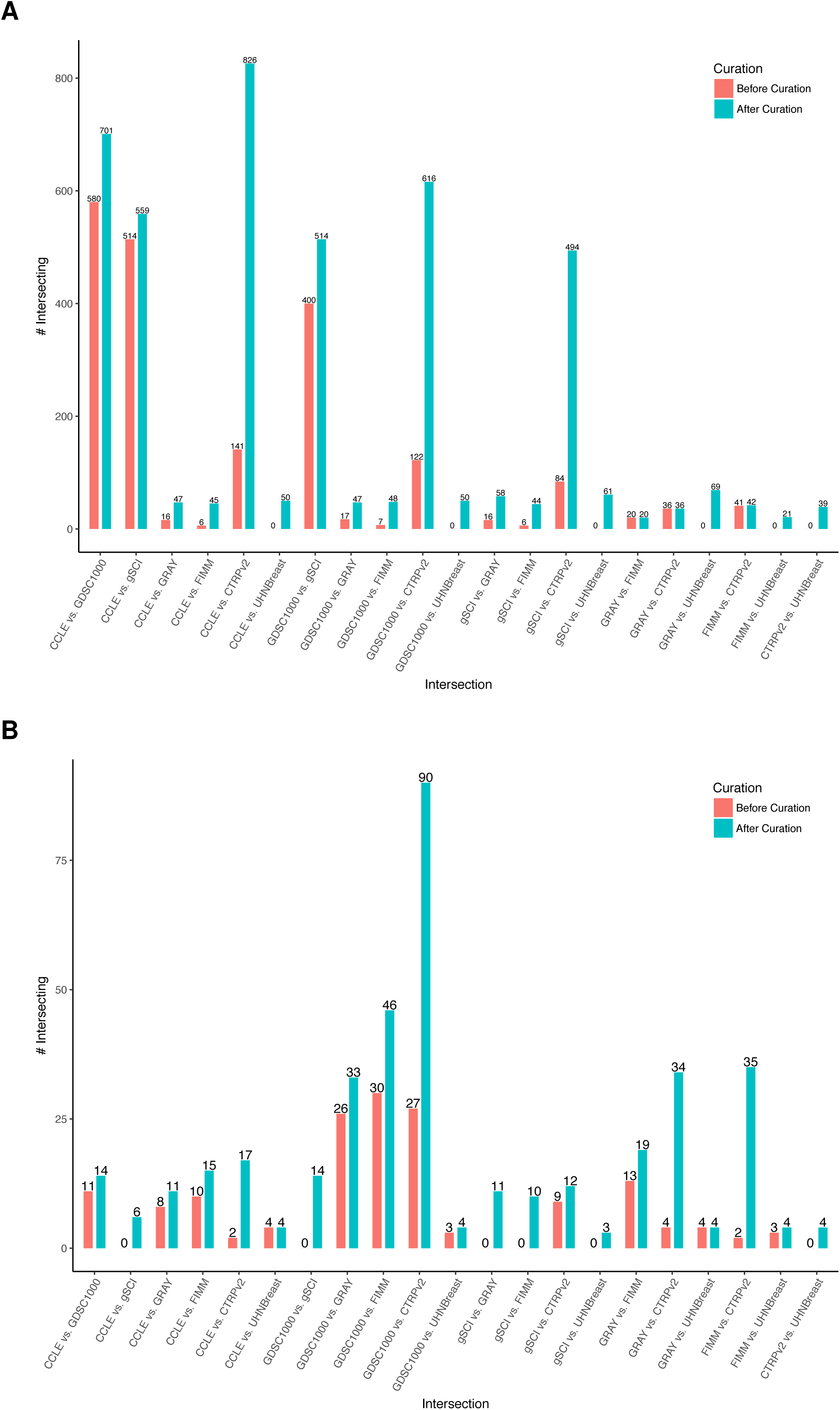
The difference in overlap between datasets before and after curation for (**A**) cell lines and (**B**) compounds.

### Annotation of drug targets

To obtain a comprehensive collection of target proteins for the compounds included in PharmacoDB, the union of known drug-target associations from four distinct data sources was integrated into the database. The CTRPv2 study released curated annotations of the protein targets for compounds (19). Additional drug target annotations were retrieved programmatically from the Drug Repurposing Hub (35), DrugBank (36), and ChEMBL (37). For DrugBank, we retrieved the gene symbol for each target using *UniProt.ws* (*v*ersion 2.16.0). For ChEMBL, we used the Web API to retrieve gene symbols for the protein targets, subsequently linked to the appropriate GeneCard (38, 39). As expected, we found tyrosine kinases to be the targets with the most associated compounds: KDR (also known as the vascular endothelial growth factor receptor 2, VEGFR-2), KIT, FLT1/3/4, and PDGFRA/B (Figure 3B). These genes are the targets of multiple kinase inhibitors, including linifanib, foretinib, tivozanib and axitinib that have been extensively tested across datasets (over 1,800 drug dose-response curves for each drug). HDAC, CDK and PI3K are also the most prevalent targets in PharmacoDB (Figure 3B).

## DATABASE ORGANIZATION AND WEB-INTERFACE

### Database implementation

All of the data is stored in a MySQL database running the default MyISAM database engine and with indexing configured on all tables in order to speed up queries (database schema in Supplementary Figure 2). The web interface is implemented using Ruby (version 2.4.1) and Ruby on Rails (version 5). To provide a smooth navigation experience, the front-end is rendered on the server and performance is optimized with use of Turbolinks (version 5.0), which does selective updates and contributes to faster page load times. All charts were produced using d3.js (version 3), a JavaScript library tailored to produce dynamic and interactive data visualizations using SVG, HTML5, and CSS web standards. Every plot generated on PharmacoDB is available for download in the SVG vectorized graphics format and the data used to generate the plot are exportable as spreadsheets.

### Search Interface

Often, biomedical researchers interested in leveraging pharmacogenomic data are investigating a specific biological question about a given cell line, tissue, drug, or target. PharmacoDB is designed to quickly answer the question: “What Pharmacogenomic data is available for my entity of interest?”. A universal search bar interface allows users to intuitively search across all entities included in the database: datasets, tissues, cell lines, compounds and targets (Figure 1; Supplementary Figure 1). This search bar is enhanced with autocompletion, giving quick feedback to the existence of an entity in our database, and helping with correct spelling and punctuation of entries. The search bar also handles more complex queries, allowing the user to search for pairwise and three way intersections of datasets, and navigate directly to a dose-response curve by querying for a cell line and compound in combination.

### Search by synonyms

The same cell line, tissue, or compound entity is often known by several names, which are often used interchangeably in the literature. As described above, semi-manual curation of datasets in PharmacoDB was done to map the synonyms used in each dataset to a unique human interpretable identifier. However, as each user may be more or less familiar with a specific synonym for a given compound or cell line, PharmacoDB was implemented such that it is possible to search for a compound, tissue or cell line by any of the synonyms collected in the curation process. This means that if a researcher is familiar with the name of a compound used in the CTRPv2 dataset, for example, they can use this identifier to find experiments with the same compound across other datasets. The synonyms encountered include different spelling or punctuation as well as completely different names, and enable a more natural interface with the database. Currently, there are 4,162 different synonyms used to refer to the 1,691 cell lines, 980 synonyms to refer to the 759 compounds, and 184 synonyms for the 41 tissues in PharmacoDB.

### Explore Interface

Complementing the Search interface, the explore page serves as a gentle entry point for new users attempting to navigate PharmacoDB. It facilitates discovery of content by presenting to the user all the entities aggregated in the database. User interaction with the explore page occurs in a series of filtering steps to find the entity or experiment of interest. Depending on which selections users make, unrelated annotations are filtered out until they make a selection corresponding to a single query of the database This will allows the user to quickly navigate through the large collection of entities while having a complete picture of all the targets, tissues, compounds and drugs included in PharmacoDB.

### Profile Pages

If a search query for a single entity is entered into the search bar, or if an entity is selected in the explore page, the user is redirected to a profile page. This page is designed to provide the user with a comprehensive view of all the data available for the entity of interest (Figure 5). Textual information is consistently positioned on the left side, and visualizations of the data are on the right. Each page contains a card in the top left corner describing the entity selected, and any relevant metadata available for this entity. For example, a cell line card would contain the relevant annotations and a description of the available molecular and pharmacological data collected for each dataset, the tissue and disease type for a cell line (Figure 5A). The card will also contain links to any relevant external databases, such as GeneCards for targets, Pubchem for compounds, and Cellosaurus for cell lines. Searchable tables list all the dose response experiments in PharmacoDB pertaining to the entity being profiled (Figure 5B). Elements in these tables tables are fully linked to allow users to navigate between profile pages in PharmacoDB, or by clicking on the experiment count for a compound and cell line pair, redirected directly to page displaying the drug testing experiment(s). The right-hand side of the profile pages displays plots with summary statistics about the entity (Figure 5C). For a cell line, a waterfall plot reports the 15 compounds with the lowest and highest efficacy or potency (see *drug dose-response curve*). These plots allow finding the most effective drug for a cell line of interest, or finding cell lines which are abnormally sensitive for a drug of interest, across all the datasets included in the database.

**Figure 5.**
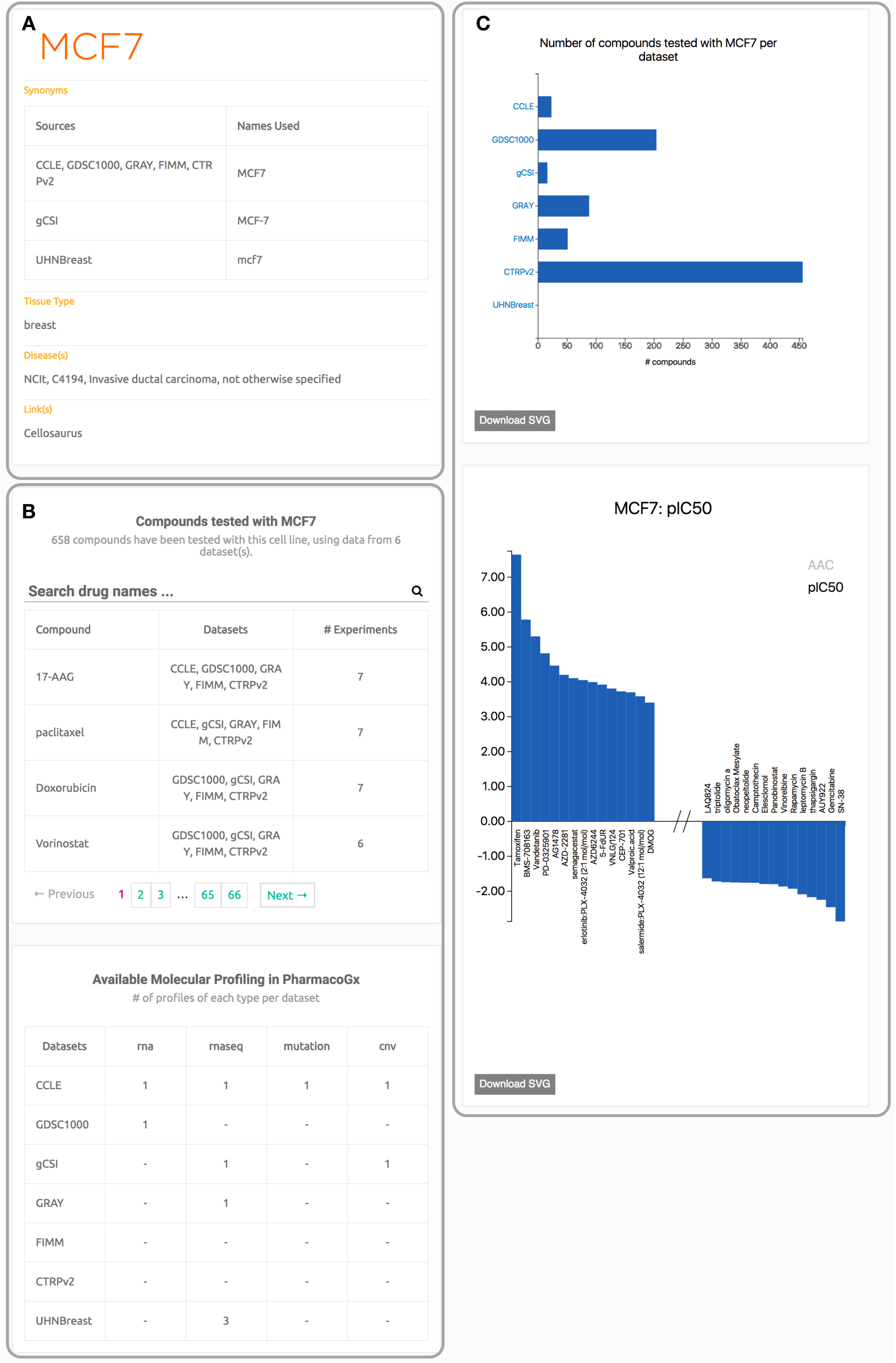
An example profile page from PharmacoDB for a cell line. The page is organized so that the left column contains textual information, and the right column contains plots. Panel (**A**) is the information card for the cell line (MCF7). Panel (**B**) contains tables listing the available data profiles for this cell. Panel (**C**) contains summary plots about the drug screening performed in each dataset and the waterfall plots of the cell response to treatment with compounds.

### Molecular data

In addition to cell line viability screens, the pharmacogenomic datasets included in PharmacoDB include extensive molecular profiling. We recently released the *PharmacoGx* package (31) to facilitate the analysis of the relationships between the pharmacological and molecular data for the purposes of biomarker discovery (30) and drug repurposing (20). The reprocessing of pharmacological data and the extensive curation of identifiers done for PharmacoDB has been fully integrated into PharmacoSet (PSet) R objects released with the *PharmacoGx* platform. While PharmacoDB does not contain molecular data, a PSet object has been created and linked to from each dataset profile page (Supplementary Figure 1A). To facilitate finding molecular data for a specific cell line, a table describing the availability of molecular profiles in *PharmacoGx* is available at the bottom of each cell line profile page. The link between PharmacoDB and *PharmacoGx* enables bioinformaticians to use PharmacoDB as an entry point for their pharmacogenomic analysis, and allows them to leverage the extensive curation done in PharmacoDB.

### Drug dose-response curves

Searching for a cell line-drug pair or selecting a cell line with a drug through the explore page will redirect to a page displaying the dose-response data found across all datasets. The page includes a plot of the measured viability values and a Hill Slope curve fit to the measured data (Figure 6A; Supplementary Methods), followed by a table of summary statistics commonly used to summarize the sensitivity/resistance of the cell line to the given compound (Figure 6B). Each curve plotted on the graph can be hidden and shown by clicking on its entry in the legend. We used *PharmacoGx* (31) to normalize and reprocess all cell viability data with a uniform pipeline to remove any biases between datasets introduced by computational aspects such as choice of Hill Slope model, curve fitting algorithms, or inconsistent calculations of summary statistics between studies (Supplementary Methods). Given the lack of consensus regarding the best way to summarize drug dose-response curve, we computed a compendium of summary metrics for the response of the cell line to the treatment with the compound, including the common IC_50_ (dose of 50% inhibition of cell viability), EC_50_ (dose at which 50% of the maximum response is observed), Area Above Curve (AAC), E_inf_ (maximum theoretical inhibition), and the recent drug sensitivity score (DSS) (40). Hovering over a value in the summary measures table will display on the plot a visualization of the procedure used to calculate the summary statistic. As there is no consensus as to the optimal metric for summarizing the information contained in the dose response curve (41). The IC_50_ and EC_50_ metrics focus on the potency of the compound, the E_max_ on the efficacy, and the AAC and DSS integrate both potency and efficacy. These metrics are presented together on PharmacoDB, and a visualization of method of calculating them is displayed directly on the drug dose-response curve to aid in making an informed decision of the correct measure to use for a given experiment or specific biological question of interest.

**Figure 6.**
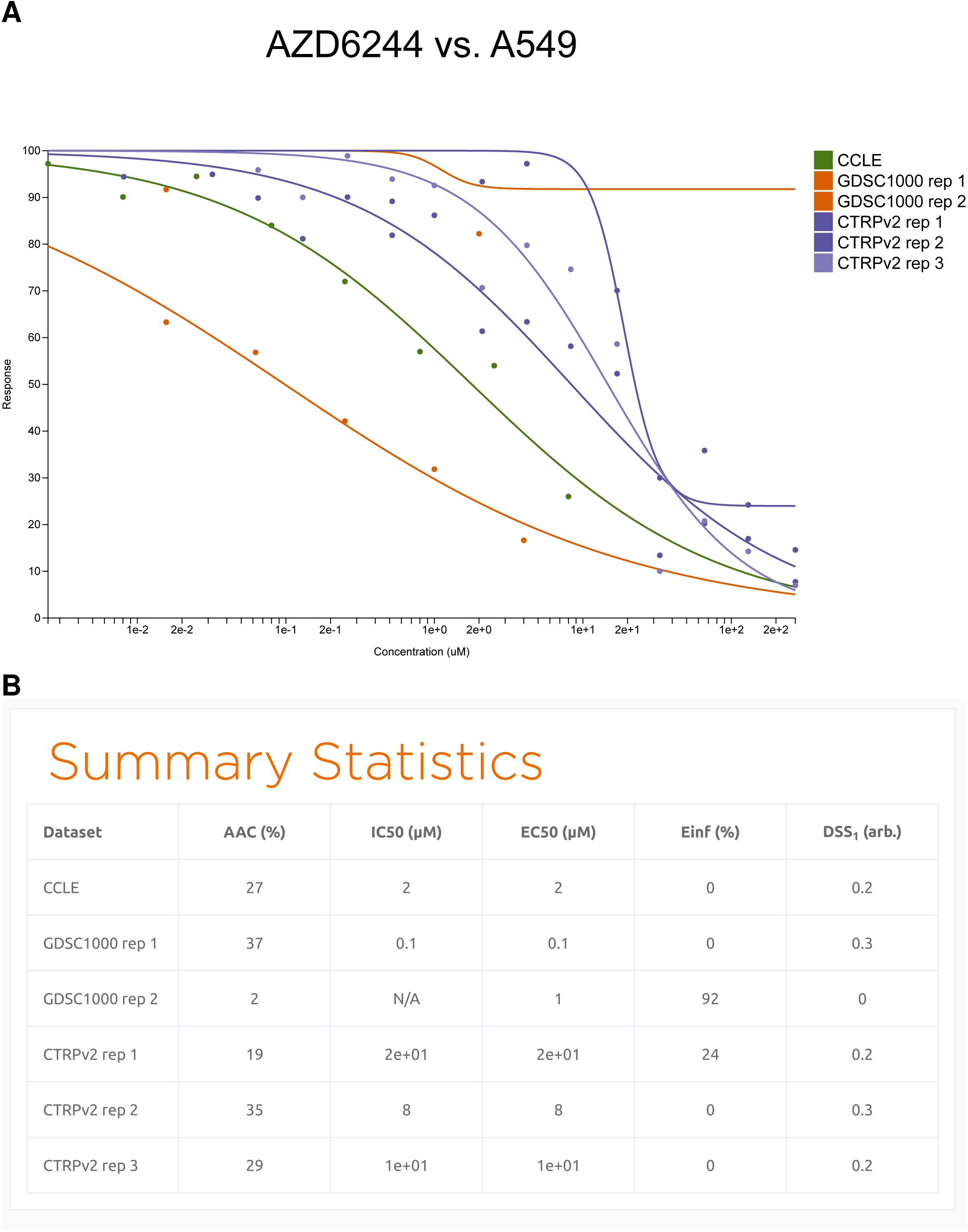
An example of drug dose response curve plot with (**A**) the A549 lung cancer cell line treated with the MEK inhibitor AZD6244, and (**B**) the corresponding table of summary statistics. IC_50_: dose of 50% inhibition of cell viability; EC_50_: dose at which 50% of the maximum response is observed; AAC: Area Above Curve; E_inf_: maximum theoretical inhibition; DSS: drug sensitivity score.

**Figure 7.**
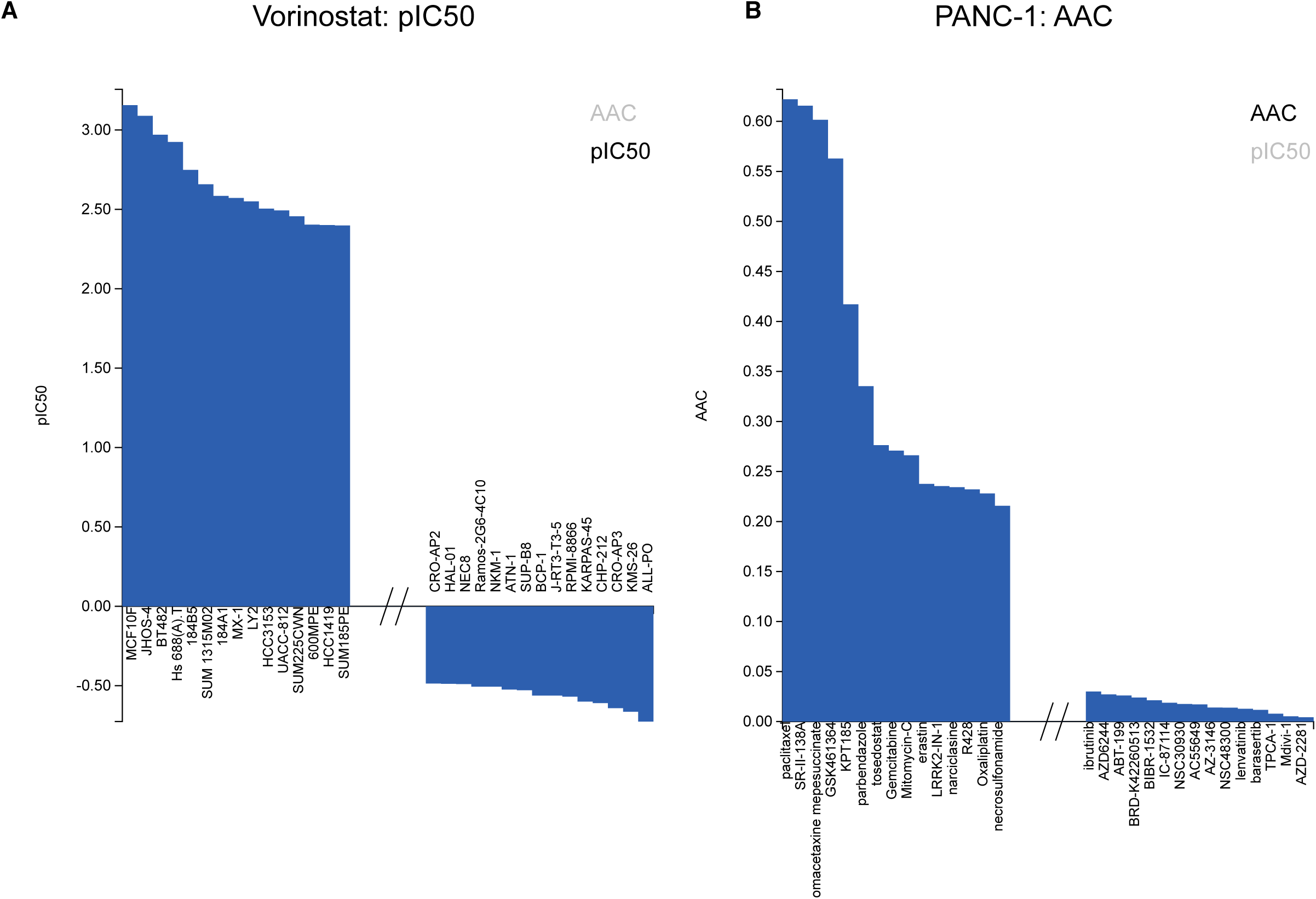
Waterfall plots of (**A**) the summarized sensitivity values across drugs for a given cell line (PANC-1) using the Area Above Curve (AAC) summary statistic and (**B**) across cell lines for a given drug (Vorinostat) using the pIC_50_ (negative logarithm base 10 of the 50% inhibitory concentration) summary statistic.

## USER ACCESS TO DATA, CODE AND FEEDBACK

### Programmatic data access

PharmacoDB exposes its data through an Application Programming Interface (API), enabling users to programmatically interact with the application. The API is RESTful (Representational State Transfer), meaning that all application resources are made available using a predefined set of stateless operations, in this case being HTTP verbs such as GET, POST, DELETE. No authentication keys, or tokens, are currently needed in order to access the API. The API has been implemented using the Go programming language, and Gin HTTP web framework has been used for routing. All queries are made using HTTP GET requests, and all results are returned in JSON format by default. The base URL used for all queries is https://api.pharmacodb.com/v1/. The end “/v1” corresponds to the version of the API being queried. This version can be changed to any of the API versions that are, or will be, released by PharmacoDB. Whenever a newer version is released, both the API and the MySQL database of the previous version are frozen and open for use at that version URL. Furthermore, breaking changes will only be introduced in a newer version. Hence, no application that integrates or uses PharmacoDB data will break, or be affected by breaking changes. All the data in PharmacoDB are publicly available via the API. Additionally, a dump of the SQL database is available for download from the front page and R objects for all the pharmacogenomic datasets are available via the *PharmacoGx* R/Bioconductor package (31).

### Code and documentation

The PharmacoDB code is open-source and publicly available through the PharmacoDB GitHub repository (github.com/bhklab/PharmacoDB) under the GPLv3 license. The documentation is available in the web-application as video and textual descriptions of all the entities and search queries in PharmacoDB. These include descriptions of the dataset, tissue, cell line, target, and drug/compound pages. Tutorial on how to perform more complex queries, such as displaying a drug dose-response curve and intersecting two or three datasets, is also described in details in the Documentation page of PharmacoDB.

### Feedback

Our web-application provides an easily accessible, optionally anonymous contact mechanism for providing feedback on all aspects of PharmacoDB. Users can suggest corrections to annotations by clicking on the feedback icon accessible on the left of every profile page. The fields in the “Contact Us” page are then prefilled with data relevant to the annotation in question. The GitHub API is then used to automatically file user suggestions and feedback as issues in the GitHub Issue Tracker at our repository (github.com/bhklab/PharmacoDB). This allows for full transparency regarding the reliability of data in the database and enables the community to fully assess and correct any missing information.

## SUMMARY AND FUTURE DIRECTIONS

PharmacoDB is the first database providing a comprehensive resource to search and explore the largest pharmacogenomic studies released to date. By combining rigorous curation of identifiers across the published pharmacogenomic datasets with comprehensive search and visualizations of the pharmacological data, PharmacoDB allows researchers to quickly access the data available to answer their biological questions of interest. It provides an interface to query for specific drug dose-response curves, and easily find the largest possible intersection between datasets.

As current pharmacogenomic datasets continue to expand and new ones are published, the number of cell lines screened with compounds will increase, opening new avenues of research for meta-analysis in biomarker discovery and other applications. In this setting, PharmacoDB will provide a unique resource where researchers can quickly mine the large amount of data generated by these high-throughput drug screening studies. Moreover, given the recent activity in the pharmacogenomic field, new statistical approaches are being developed to better model and summarize drug dose-response curves. Recently, Hafner et al. published the growth rate inhibition 50 (GR_50_) metric to robustly quantify drug response by accounting for the different proliferation rate of each cancer cell lines (42), and showed an increase in consistency across datasets (43). Although this method and others may require data that are not always available for all datasets (e.g., proliferation rate of each cell lines for GR_50_) we are committed to implement them to provide users with the opportunity to select the most relevant readout for their specific application. In addition to datasets measuring cell viability, we also plan to update PharmacoDB with pharmacogenomic datasets reporting the transcriptional changes due to chemical perturbation, such as the Connectivity Map (17) and the L1000 (44) datasets. The combination of drug sensitivity and perturbation data would allow users to study deeper the relationship between the molecular state of cancer cell lines and their response to compound perturbations (20). The flexibility of PharmacoDB will enable continuous update of the pharmacogenomic datasets, and facilitate the analysis of these valuable data by the scientific community.

## SUPPLEMENTARY DATA

**Supplementary Methods** reporting the sources of pharmacogenomic datasets, and describing the fitting of the drug dose-response curves and the application programming interface (API).

**List of acronyms** used throughout the manuscript.

**Supplementary Figure 1:**
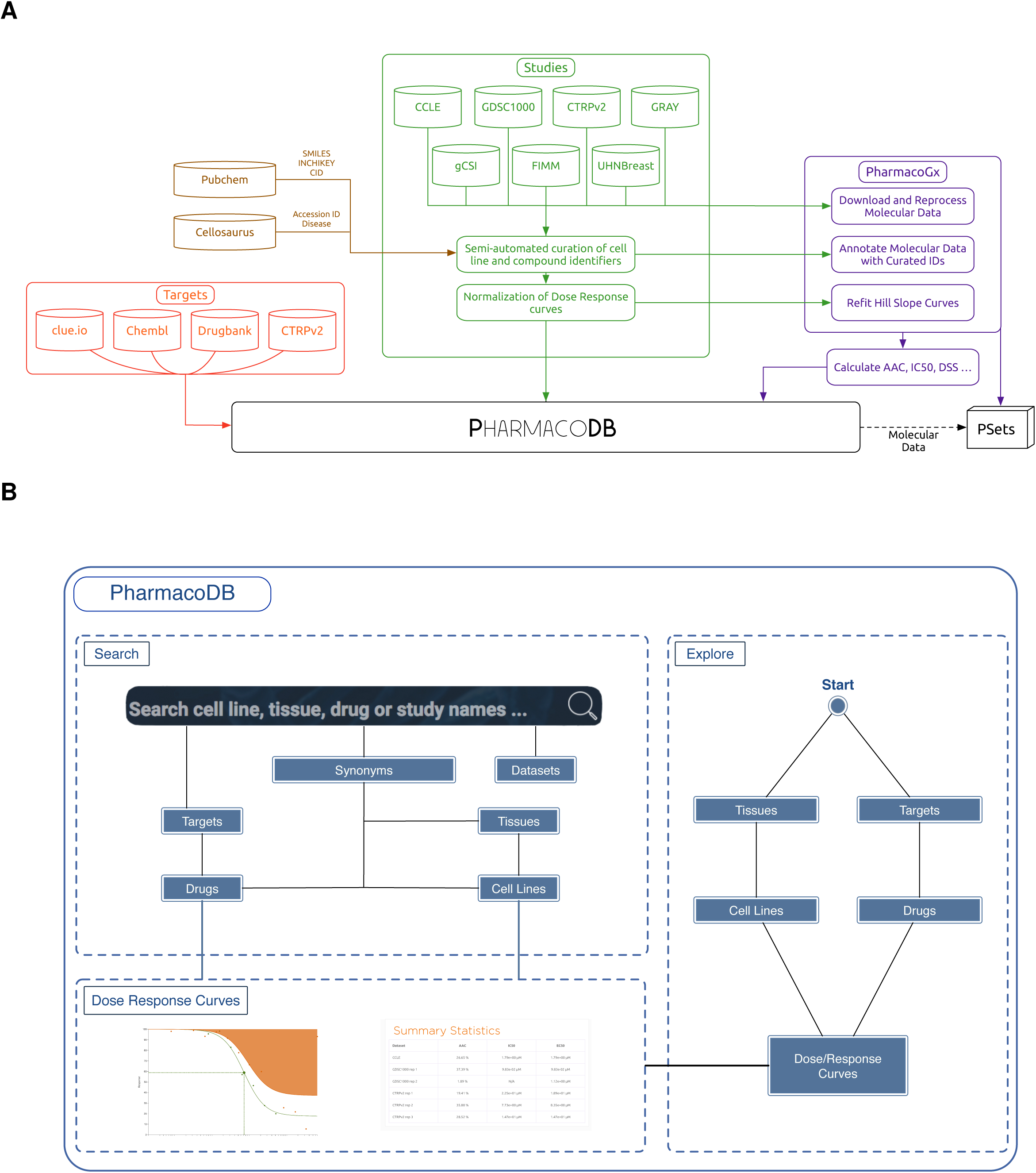
Flow diagrams describing (**A**) the external resources used to populate PharmacoDB; and (**B**) the “Search” and “Explore” interfaces to query PharmacoDB.

**Supplementary Figure 2:**
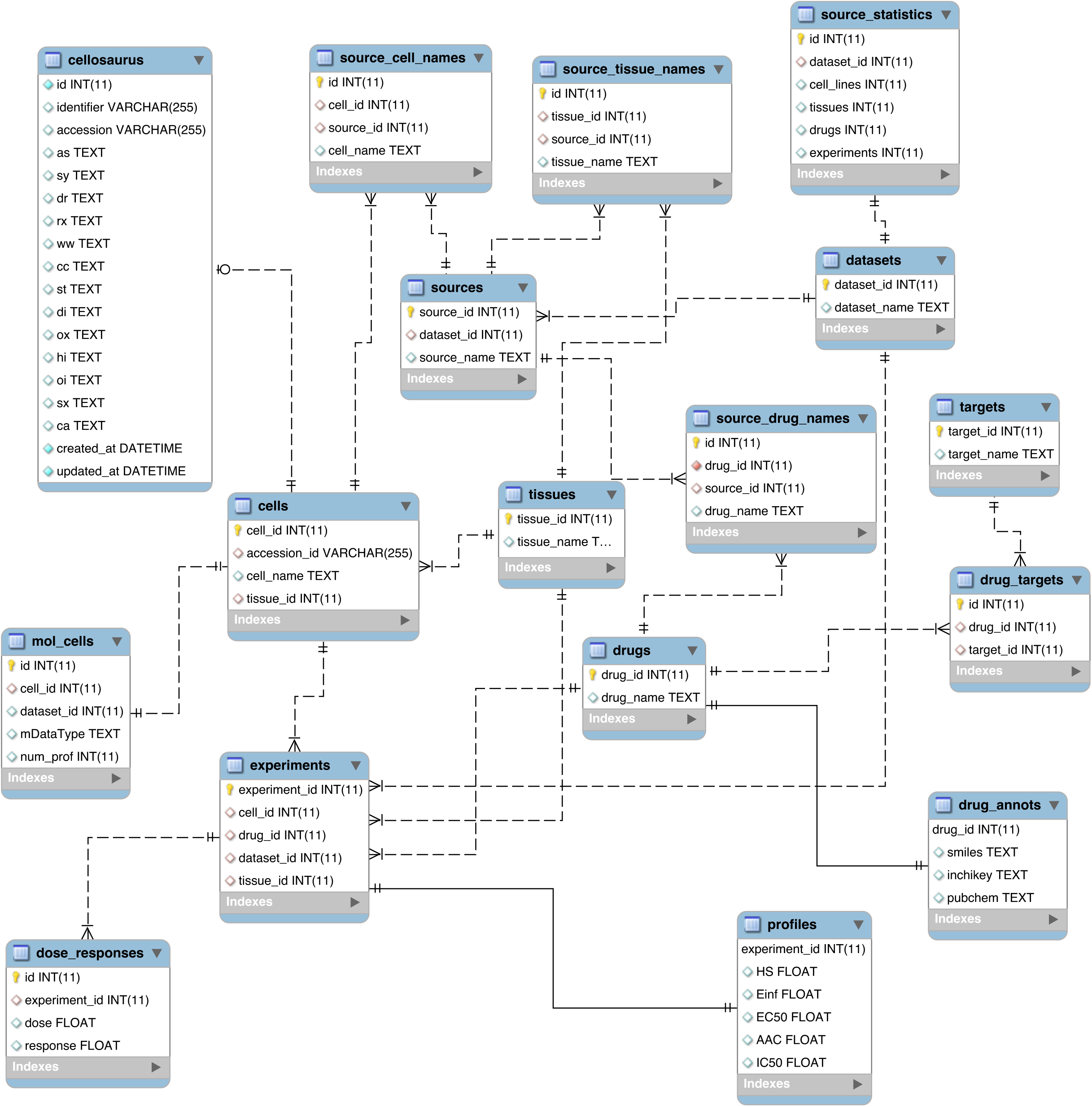
Structure of the database at the level of SQL tables and their relationships.

## ACKNOWLEDGEMENTS

The authors would like to thank the investigators of the Genomics of Drug Sensitivity in Cancer (GDSC), the Cancer Cell Line Encyclopedia (CCLE), Genentech (gCSI), the Institute for Molecular Medicine Finland (FIMM), Drs. Joe W. Gray, Benjamin G. Neel and David W. Cescon who have made their valuable pharmacogenomic data available to the scientific community. The authors also thank Heather Selby for her help curating the cell line identifiers, and Aimos Bairoch for developing Cellosaurus and helping us obtain detailed annotations for our collection of cell lines.

## SUPPLEMENTARY INFORMATION

### Supplementary Methods

#### Sources for the pharmacogenomic studies

For the Cancer Cell Line Encyclopedia, the data was downloaded from the CCLE portal (https://portals.broadinstitute.org/ccle), using the February 24, 2015 update of the pharmacological profiling data. The data from the Genomics of Drug Sensitivity in Cancer project was downloaded from the dedicated portal (http://www.cancerrxgene.org/), using release 6 of the data, corresponding to the GDSC1000 update of the dataset. The gCSI dataset was made available with Haverty et al. (Nature 2016) in the compareDrugScreens R package (http://research-pub.gene.com/gCSI-cellline-data/), which was fetched July 2016. The CTRP dataset was retrieved from the National Cancer Institute Cancer Target Discovery and Development (CTD2) data portal, using the v2 release of the data (CTRPv2; https://ocg.cancer.gov/programs/ctd2/data-portal). The OHSU GRAY dataset was obtained from the supplementary data of Daemen et al. (Genome Biology 2013).

#### Fitting drug dose-response curves

The rate-limiting step in the killing of susceptible cancer cells by an anti-cancer drug is assumed to be the binding of a target receptor *R* to *n* molecules of a drug *D*, and the free drug molecules and unbound target are assumed to be in thermodynamic equilibrium with the bound drug-target complex:

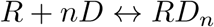

By the Law of Mass Action, this equilibrium is characterized by an equilibrium constant *K* satisfying the equation

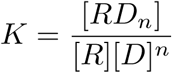

where brackets around a variable name denote the concentration of the molecule it represents. It then follows that the fraction *f* of receptors bound to ligands is given by

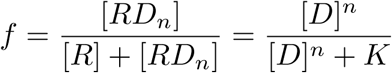

Since this binding is the rate-limiting step in the killing of cancer cells, the fraction *y*(*x*) of susceptible cancer cells killed by a concentration *x* of the drug is approximately equal to the fraction of unbound receptors 1 – *f* :

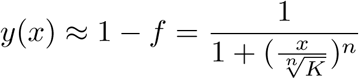

We define 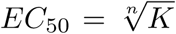 and interpret it as the concentration of the drug needed to have half of the target receptors bound at equilibrium. We also assume that a fraction *E*_*∞*_ of the cancer cells are not susceptible to the drug at all. This leads to our final equation

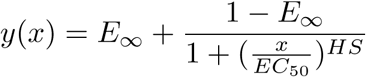

where *y*(*x*) = 0 denotes death of all treated cells, *y*(*x*) = 1 denotes no effect of the drug dose, *EC*_50_ is the concentration at which 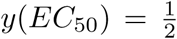, and *HS* is a parameter describing the cooperativity of binding. *HS <* 1 denotes negative binding cooperativity, *HS* = 1 denotes noncooperative binding, and *HS* > 1 denotes positive binding cooperativity.

This is the basic mathematical structure that was posited to underlie the dose-response data observed in the study. Consequently, median cellular viability data from all datasets was fit by means of least-squares regression to equations of this type. To ensure robustness of the curve-fitting algorithms, bounds were placed on the values of each of these parameters. Drugs were assumed not to increase the fitness of malignant cells, so *E*_*∞*_ was constrained to lie in the interval [0, 1]. Drugs were also assumed to have *EC*_50_ values within [1pM, 1M], an interval containing the *EC*_50_ values reported by Barretina et al. (Nature 2012). Finally, we follow Fallahi et al. (Nat Chem Biol 2013) in allowing HS to lie anywhere in [0, 4].

Barretina et al. (Nature 2012) fit dose-response data to one of three models. In most cases, their model of choice was identical to our own, with the addition of a maximum viability parameter *E*_0_. Their dose response equation then became

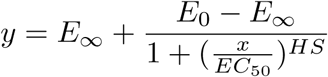

The inclusion of this parameter makes comparison of dose-response curves problematic. With its inclusion, the viability of the cell line in the absence of any drug becomes

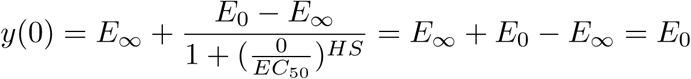

As a result, the viability measures of different drug-cell line combinations are normalized differently, and direct comparison of viability predictions from different dose-response curves is no longer appropriate. The *IC*_50_ values they reported, however, were simply the concentrations at which their fitted curves reached viability reduction of 50% of cellular viability. The end result was a reported *IC*_50_ value that assumed normalization of viability data to the negative control associated with a curve fitted assuming normalization of viability data to a reference level that was most consistent with the observed data. The *IC*_50_ values published in the paper’s supplementary information thus represented viability reduction by a fraction that varied from cell line to cell line.

In GDSC, the following five-parameter model was used:

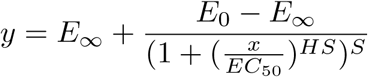

However, since the *E*_0_ parameter is fixed by controls, their curve can be represented as

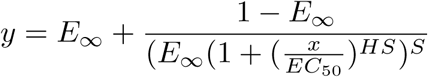

This parameter accounts for the presence of an antagonistic binding of the drug, and introduces asymmetry into the theoretical log dose-response curve. The extra parameter, known as the “Schild slope”, allows the dose-response curve to be non-monotonic.

While this parameter is well-founded biologically, we chose not to use it in our own dose-response curves. As only medians of technical replicates are available for CCLE, using a 4-parameter model would have increased our susceptibility to overfitting noise in the sparse dose-response curves. Furthermore, we only rarely observed the non-monotonicity that necessitates the inclusion of a Schild slope parameter in a very small fraction of dose-response curves. For these reasons, we ultimately chose to use our simpler 3-parameter model to compare the dose-response curves from the GDSC and CCLE datasets.

### Application Programming Interface (API)

#### Overview

The API currently supports a wide range of search queries, based on the resource type being queried. Resource types include: *cell lines*, *tissues*, *drugs*, *datasets*, *experiments*, *intersections*, and *stats*. All available endpoints for each resource type are documented in the API’s main repository on github at https://github.com/bhklab/PharmacoDB.

#### Request/Response Format

All requests use HTTP GET method, and the base URL used for making requests is https://api.pharmacodb.com/v1/. Valid endpoints should be concatenated to the base URL when making a request. All valid endpoints are listed on github, as well as the **API Reference** section below. No API keys, or tokens, are needed in order to make an API call.

To demonstrate, the following is a sample request using the **curl** command:

$ curl “https://api.pharmacodb.com/v1/cell_lines”

The above request queries a list of cell lines found in PharmacoDB. All resource requests return a paginated list by default. This means that if PharmacoDB contans N number of cell lines, only the first K items are returned at initial request, and the rest N - K items can be queried by changing the page value. The *page* and *per_page* options are available for modifying paginated lists. For example, the above request can be modified to get the third page, whereby each page contains 20 items, as follows:

$ curl “https://api.pharmacodb.com/v1/cell_lines?page=3&per_page=20”

If, instead, all cell lines need to be queried in a single request, the request will look as follows:

$ curl “https://api.pharmacodb.com/v1/cell_lines?all=true”

Setting the *all=true* option overrides default pagination, and returns a list of all items found in a resource type of interest.

To select a single item in a resource type, an ID is used as a parameter, as follows:

$ curl “https://api.pharmacodb.com/v1/cell_lines/{id}”

In the above, *id* corresponds to either a cell line ID, or a cell line name. By default, the API assumes the request is using a cell line ID. For example, making a request to:

$ curl “https://api.pharmacodb.com/v1/cell_lines/1”

retrieves the cell line whose ID = 1. In order to query a cell line by its name instead of ID, one can use the *type* option, as follows:

$ curl “https://api.pharmacodb.com/v1/cell_lines/mcf7?type=name”

Metadata, such as pagination information and links, are all included in response header by default. This metadata can be included in response body by using the *include=metadata* option. For example, making a request to:

$ curl “https://api.pharmacodb.com/v1/cell_lines?include=metadata”

retrieves a list of cell line items, whereby pagination information and links metadata are included in response body.

All results are indented, or pretty printed, by default. Users can customize results and turn off indentation by using the *indent=false* option, as follows:

$ curl “https://api.pharmacodb.com/v1/cell_lines?indent=false”

All endpoint options are listed below in the **API Reference** section.

#### Errors

All valid responses return with HTTP status code 200 (Status OK). Any other status code in response signals an error. There are three main error types defined in the API.

1. **400 (Bad Request)**: This error type is returned when no routers match the request URL. This can happen when the requested endpoint is not one that is defined in the API. The API documentation lists all available endpoints.
2. **404 (Not Found)**: A 404 error is returned whenever a resource item is not found in database.
3. **500 (Internal Server Error)**: This is returned whenever a general error occurs in the API itself instead of user query. Users should contact us if met with 500 Internal Server Error responses.

#### API Reference

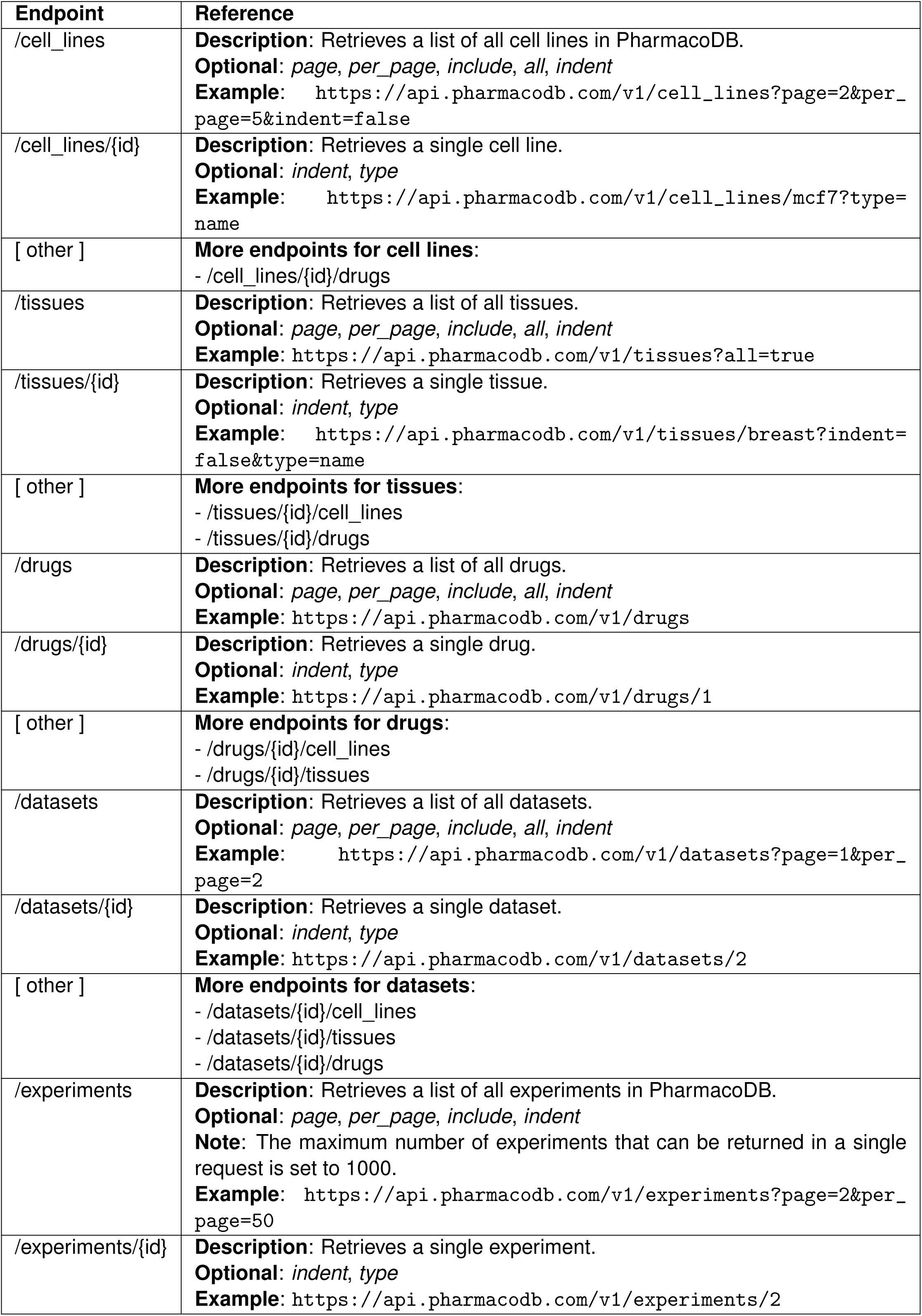
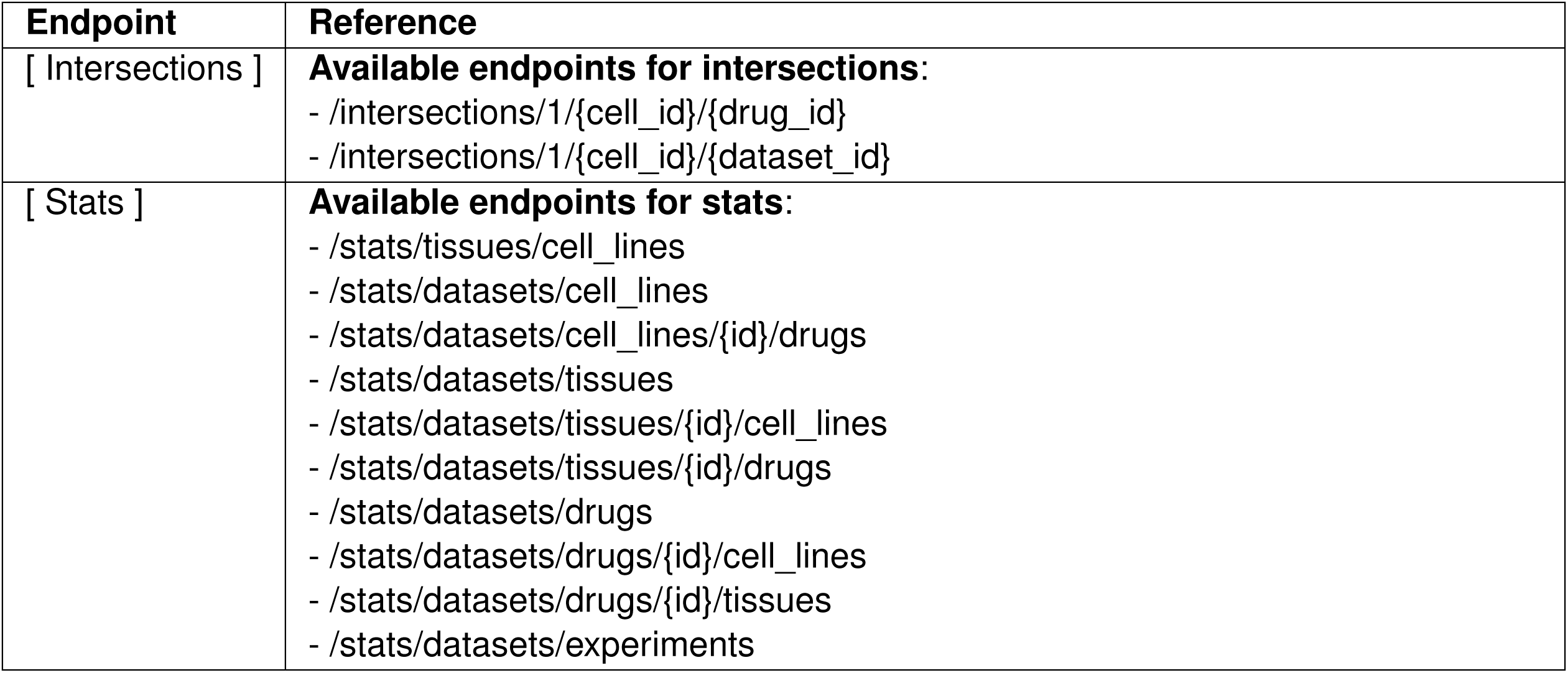

### Acronyms

AAC: Area above the dose response curve
CCLE: The Cancer Cell Line Encyclopedia initiated by the Broad Institute of MIT and Harvard
CMAP: Connectivity Map by the Broad Institute
CTRP: Cancer Therapeutic Response Portal
DSS: Drug sensitivity score
EC_50_: Dose at which 50% of the maximum response is observed
E_*max*_: Maximum theoretical inhibition
FIMM: Institute for Molecular Medicine Finland cell viability screen
gCSI: Genentech Cell Screening Initiative
GDSC: The Cancer Genome Project initiated by the Wellcome Trust Sanger Institute
IC_50_: Concentration at which the drug inhibited 50% of the maximum cellular growth
InchiKey: International Chemical Identifier
OHSU: Oregon Health and Science University
pIC_50_: –log_10_(IC_50_)
SMILES: Simplified molecular-input line-entry system
UHN: University Health Network

